# Metabolic influence of core ciliates within the rumen microbiome

**DOI:** 10.1101/2022.06.22.497163

**Authors:** Thea O. Andersen, Ianina Altshuler, Arturo V.P. de Leon, Juline Walter, Emily McGovern, Kate Keogh, Cécile Martin, Laurence Bernard, Diego P. Morgavi, Tansol Park, Zongjun Li, Yu Jiang, Jeffrey L. Firkins, Zhongtang Yu, Torgeir R. Hvidsten, Sinead M. Waters, Milka Popova, Magnus Ø. Arntzen, Live H. Hagen, Phillip B. Pope

## Abstract

Protozoa comprise a major fraction of the microbial biomass in the rumen microbiome, of which the entodiniomorphs (order: *Entodiniomorphida*) and holotrichs (order: *Vestibuliferida*) are consistently observed to be dominant across a diverse genetic and geographical range of ruminant hosts. Despite the apparent core role that protozoal species exert, their major biological and metabolic contributions to rumen function remain largely undescribed *in vivo*. Here, we have leveraged (meta)genome-centric metaproteomes from rumen fluid samples originating from both cattle and goats fed diets with varying inclusion levels of lipids and starch, to detail the specific metabolic niches that protozoa occupy in the context of their microbial co-habitants. Initial proteome estimations via total protein counts and label-free quantification highlight that entodiniomorph species *Entodinium* and *Epidinium* as well as the holotrichs *Dasytricha* and *Isotricha* comprise an extensive fraction of the total rumen metaproteome. Proteomic detection of protozoal metabolism such as hydrogenases (*Dasytricha, Isotricha, Epidinium, Enoploplastron*), carbohydrate-active enzymes (*Epidinium, Diplodinium, Enoploplastron, Polyplastron*), microbial predation (*Entodinium*) and volatile fatty acid production (*Entodinium* and *Epidinium*) was observed at increased levels in high methane-emitting animals. Despite certain protozoal species having well-established reputations for digesting starch, they were unexpectedly less detectable in low methane emitting-animals fed high starch diets, which were instead dominated by propionate/succinate-producing bacterial populations suspected of being resistant to predation irrespective of host. Finally, we reaffirmed our abovementioned observations in geographically independent datasets, thus illuminating the substantial metabolic influence that under-explored eukaryotic populations have in the rumen, with greater implications for both digestion and methane metabolism.

## BACKGROUND

Ruminants operate in symbiosis with their intrinsic rumen microbiome, which is responsible for the degradation of forage into nutrients, in the form of volatile fatty acids (VFAs), supplying ∼70% of net energy for the host^1^. The rumen microbiome itself is a complex assemblage of bacterial, fungal, archaeal, viral, and protozoal microorganisms whose intricate composition and function is connected to host productivity traits, such as feed efficiency, milk yield, animal health, and greenhouse gas (GHG) emissions^2-5^. Large collaborative research efforts have been made to identify and characterize the core rumen microbiome including creating a publicly available catalogue for cultivated and sequenced genomes^6-8^. In the rumen, bacteria are estimated to constitute 50-90 %, protozoa 10-50 %, fungi 5-10% and archaea less than 4% of the total microbial biomass^9,10^. Due to the difficulties of axenically culturing rumen eukaryotic populations and their complex genomic features that are obstinate to current metagenomic assembly and binning technologies, the reconstruction of the rumen microbiome has been heavily biased towards bacterial and archaeal members, whereas the fungal and protozoal contributions of the rumen currently remain poorly characterized. While anaerobic fungi have a reputable role as fibre degraders in the rumen, only 18 anaerobic gut fungi from herbivores are currently described, with only 11 genomes available^11-13^. Similarly, to date few rumen protozoal genomes are sequenced and publicly available, chief among them, the rumen ciliate protozoa *Entodinium caudatum*^14^. More recently, single-cell amplified genomes (SAGs) have been recovered from rumen microbiome samples representative of 5 holotrich and 14 entodiniomorph species spanning 13 genera including *Isotricha, Dasytricha, Diplodinum, Enoploplastron, Metadinium, Eremoplastron, Ostracodinium, Polyplastron, Ophryoscolex, Epidinum* and *Entodinium*^15^.

*Entodinium* and *Epidinium* are entodiniomorphs of the order *Entodiniomorphida* and represent two of the most dominant genera of rumen protozoa, previously being detected in more than 99% of 592 rumen samples at a mean protozoal relative abundance of ∼ 38% and 16% respectively (2015 rumen census: 32 animal species, 35 countries)^16^. Entodiniomorphs have been previously observed to stimulate methane production^17^, but while some possess hydrogenosomes such as *Epidinium caudatum*^18^, it is believed the numerically dominant *Entodinium caudatum* does not, and instead encodes mitosomes and iron hydrogenases that may indicate hydrogen production, beneficial for methanogenic endosymbionts^19^. Holotrich species of the order *Vestibuliferida* such as *Dasytricha* and *Isotricha* are generally found at lower abundances compared to entodiniomorphs but are renowned for their fermentation of soluble carbohydrates and their production of VFAs and hydrogen^20,21^. Here, we present a genome-centric metaproteomics analysis of the rumen microbiome from two different host species; Holstein dairy cattle (*Bos taurus*) and alpine goats (*Capra hircus*), that were fed diets of first-cut grassland hay with a 45:55 forage:concentrate ratio, with concentrates supplemented with either no additional lipid (CTL), or corn oil and wheat starch (COS)^5,22,23^. Previous microscopy analysis revealed high counts of entodiniomorphs and to a lesser extent holotrichs, while metadata revealed that animals fed COS displayed reduced methane emissions, irrespective of host. To describe how these diets affect digestion and production of methane and VFAs, we sought to investigate changes in function and composition in the complex rumen microbiome. By using shotgun metagenomic sequencing we recovered in total 244 prokaryote metagenome-assembled genomes (MAGs) that together with selected isolate- and SAG-derived eukaryote genomes^14,15,24-29^ formed the database for our genome-centric metaproteomic analysis of rumen protozoal species. Key protozoal metabolisms were detected related to plant fiber and starch degradation, bacterial predation, and hydrogen and VFA production. Despite numerous protozoal species having the genetic ability to degrade starch, our analysis showed contrasting data that suggests rumen protozoa were less metabolically active in the rumen microbiome of animals fed a starch-rich diet. In such a scenario other starch-degrading and/or propionate and succinate producing bacterial genera, as *Prevotella* and *Fibrobacter* and members of the families *Succinivibrionaceae* and *Aminobacteriaceae* appeared to be more prevalent. In concert, our analysis showed that reduced methane production in both cattle and goats eating feeds supplemented with COS is likely caused via a redirection of hydrogen to succinate and propionate production instead of methanogenesis. Finally, by analysing a secondary, geographically independent dataset, we reaffirmed our primary observations of protozoal dominance and patterns concerning hydrogen, plant fiber and starch-related metabolism, thus supporting our hypothesis that this protozoal species plays a core role in rumen microbiome function.

## METHODS

### Animal trial and sample handling

The experimental procedures were approved by the Auvergne-Rhône-Alpes Ethics Committee for Experiments on Animals (France; DGRI agreement APAFIS#3277–2015121411432527 v5) and complied with the European Union Directive 2010/63/EU guidelines. Experiments were performed at the animal experimental facilities of HerbiPôle site de Theix at the Institut National de la Recherche pour l’Agriculture, l’Alimentation l’Environnement (INRAE, Saint-Genès-Champanelle, France) from February to July 2016. Experimental design, animals and diets were as described elsewhere^5,23^. Briefly, four Holstein cows and four Alpine goats, all lactating, were enrolled in respectively two 4 × 4 Latin square design trials to study the effects of 4 diets over four 28-d experimental periods (**Supplementary Table S1**). The original study included four different experimental grassland hay basal diets with concentrates supplemented with various lipid sources; control diet with no added lipids (CTL), diet supplemented with corn oil and wheat starch (COS), diet supplemented with marine algae powder (MAP) and diet supplemented with hydrogenated palm oil (HPO) (**Supplementary Table S2**)^23^. In the present study, we focused on the CTL and COS diets, which were associated with the most extreme methane (CH_4_) emission profiles in both ruminant species. The CTL diet composed of grass hay *ad libitum* with concentrates containing no additional lipid, whereas COS contained corn oil (5.0% total dry matter intake (DMI)) and wheat starch -5.0 % of total DMI (COS) (**Table 1**). Corn oil (Olvea, Saint Léonard, France) was added to the concentrate, at 5% of total DMI and contained (g/kg of total FA): 16:0 (114), 18:0 (16.4), cis-9 18:1 (297), cis-11 18:1 (6.30), 18:2n-6 (535), 18:3n-3 (7.57), 20:0 (3.48), 22:0 (1.0), 24:0 (1.5), and total FA (1000 g/kg). Detailed diet composition is available in Martin *et al*.^*5*^. Each experimental period lasted for 28 days. Diets were offered as two equal meals at 0830 and 1600h. Animals had access to a constant supply of freshwater *ad libitum*. Concentrate and hay refusals were weighed daily. The amounts of feed offered the following day was adjusted regarding to refusals to maintain the targeted the dietary 45 % dry matter (DM) forage and 55 % DM concentrate ratio. We acknowledge that by only including one sampling point for rumen fluid we are reducing the power that the original experimental Latin square design gives regarding minimizing individual error between the animals.

**Table 1.** Summary of the animal host, dietary conditions and accompanying metadata that are linked to the rumen samples used to explore protozoal functions. Effects of control diet (CTL) and diet supplemented with corn oil and wheat starch (COS) on average VFA concentration in the percentage of total VFA concentration (mmol/l) and average methane yield (CH_4_ g/kg DMI) in addition to average total protozoa cell counts and average cell counts for small entodiniomorph protozoa for cattle (n=4) and goats (n=4) (**Supplementary Table S2**)^5^. Total protozoal cell counts, and cell counts of entodiniomorphs and holotrichs were based on ciliate protozoa morphology in microscopy^5^. Measurements on VFA concentrations and methane yield were determined by gas chromatography and respiration chambers, respectively.

| Animal | Cattle |  | Goat |  |
| --- | --- | --- | --- | --- |
| Diet | CTL | COS | CTL | COS |
| VFA sum mmol/l | 61.8 | 68.3 | 33.7 | 25.3 |
| Acetate (% sum) | 72.0 | 69.1 | 65.7 | 64.2 |
| Propionate (% sum) | 14.2 | 21.3 | 15.6 | 19.1 |
| Butyrate (% sum) | 10.8 | 6.4 | 13.7 | 9.9 |
| CH<sub>4</sub> g/kg dry matter intake | 19.5 | 14.5 | 19.5 | 13.5 |
| Standard deviation | 2.2 | 2.9 | 4.0 | 4.1 |
| Total protozoa (10<sup>3</sup> cells/ml) | 92 | 100 | 1382 | 898 |
| Small entodiniomorphs (<100µm) (10<sup>3</sup> cells/ml) | 51 | 97 | 1346 | 898 |
| Large entodiniomorphs (>100µm) (10<sup>3</sup> cells/ml) | 25.1 | 1.7 | 4.9 | 0.2 |
| Holotrichs |  |  |  |  |
| *Isotricha* spp | 4.1 | 1.0 | 4.6 | 0.2 |
| *Dasytricha* spp | 11.8 | 0.1 | 26.7 | 0.2 |

Rumen fluid was collected through stomach-tubing before the morning feeding on day 27 of each experimental period. The stomach tube consisted of a flexible 23 mm external diameter PVC hose fitted to a 10 cm-strainer at the head of the probe for cattle, and a flexible 15 mm PVC hose with a 12 cm-strainer for goats. The first 200 ml of rumen fluid was discarded from to minimize contamination from saliva. Samples were filtered through a polyester monofilament fabric (280 μm pore size), dispatched in 2-ml screw-cap tubes, centrifuged at 15000 x g for 10 mins and the pellet snap-frozen in liquid nitrogen. Samples were stored at -80°C until DNA extraction using the Yu and Morrison bead-beating procedure^30^. In total, 32 rumen fluid samples (four cows and four goats fed four diets included in the original study^23^) were sent to the Norwegian University of Life Sciences (NMBU) for metagenomic and metaproteomic analysis. Respiration chambers were used to measure methane emissions over a 5-day period, while VFA and NH_3_ concentrations were determined by gas chromatography using a flame ionization detector^31^. Protozoa were counted by microscopy and categorized as either small entodiniomorphs (<100 μm), large entodiniomorphs (>100 μm) or as holotrichs *Dasytricha* and *Isotricha*^9^. Further specifics about diets and measurements can be found in Martin *et al*.^*5*^ and VFA and methane measurements are summarized in **Table 1**.

### Metagenomic sequencing and analysis

Metagenomic shotgun sequencing was performed at the Norwegian Sequencing Centre on two lanes of the HiSeq 3/4000 (Illumina) generating 150 bp paired-end reads in both lanes. Sequencing libraries were prepared using the TruSeq DNA PCR-Free High Throughput Library Prep Kit (Illumina) prior to sequencing. All 32 samples (four cows and four goats fed four diets included in the original study^23^) were run on both lanes to prevent potential lane-to-lane sequencing bias. FASTQ files were quality filtered and Illumina adapters removed using Trimmomatic^32^ (v. 0.36) with parameters -phred33 for base quality encoding, leading and trailing base threshold set to 20. Sequences with an average quality score below 15 in a 4-base sliding window were trimmed and the minimum length of reads was set to 36 bp. MEGAHIT^33^ (v.1.2.9) was used to co-assemble reads originating from samples collected from cattle and goats separately, with options –kmin-1pass, --k-list 27,37,47,57,67,77,87, --min-contig-len 1000 in accordance with^7^. Bowtie2^34^ (v. 2.3.4.1) was used to map reads back to the assemblies and SAMtools^35^ (v. 1.3.1) was used to convert SAM files to BAM format and index sorted BAM files.

The two co-assemblies (one from the samples originating from cattle and the other originating from the samples of goats) were binned using Maxbin2^36^, MetaBAT2^37^ and CONCOCT^38^. MetaBAT2 (v. 2.12.1) was run using parameters –minContig 2000 and –numThreads 4, Maxbin2 (v. 2.2.7) ran with default parameters and -thread 4, min_contig_length 2000, and CONCOCT (v. 1.1.0) ran with default parameters and –length_threshold 2000. Further, bins were filtered, dereplicated and aggregated using DASTool^39^(v. 1.1.2) with the parameters –write_bins 1, --threads 2 and BLAST^40^ as search engine. This resulted in a total of 244 dereplicated MAGs across the two host species (104 originating from cattle and 140 from goats). CheckM^43^(v. 1.1.3) lineage workflow was used to calculate completeness and contamination of each MAG, with parameters –threads 8, --extension fa, and CoverM (v. 0.5.0) (https://github.com/wwood/CoverM) was used to calculate relative abundance of each MAG, while GTDB-tk^44,45^ (v. 1.3.0) was used for taxonomic annotation. Approximately 90% (219 of 244) of the recovered MAGs were considered high or medium quality MAGs according to MIMAGs threshold for completeness and contamination for genome reporting standards^46^ (**Supplementary Table S3**). Gene calling and functional annotation of the final MAGs were performed using the DRAM^47^ pipeline with the databases dbCAN^48^, Pfam^49^, Uniref90^50^, Merops^51^, VOGdb and KOfam^52^. The translated amino acid sequences from the publicly available drafted *En. caudatum* macronucleus genome were annotated with the KEGG metabolic pathway database using BlastKOALA^53^ by Park *et al*.^14^. Proteins with resulting KEGG Orthology (KO) numbers were functionally assigned to metabolic pathways using KEGG Mapper Reconstruct Tool^54^.

The resulting protein sequences for each MAG, as well as those from the host genomes of goat (*Capra hircus*, NCBI ID: 10731) and cattle (*Bos taurus*, NCBI ID: 82) were compiled into two databases, from now on referred to as sample specific RUmen DataBase for Goat (RUDB-G) and sample specific RUmen DataBase for Cattle (RUDB-C). In addition, both databases were supplemented with the genome of the protozoa *Entodinium caudatum*^14^, as well as 18 SAGs from Li et al.^15^ including *Dasytricha ruminantium* SAG3, *Diplodinium dentatum* SAGT1, *Diplodinium flabellum* SAG1, *Enoploplastron triloricatum* SAGT1, *Entodinium bursa* SAG3, *Entodinium longinucleatum* SAG4, *Epidinium cattanei* SAG3, *Epidinium caudatum* SAG1, *Eremoplastron rostratum* SAG2, *Isotricha intestinalis* SAGT2, *Isotricha prostoma* SAG3, *Isotricha* sp YL-2021a SAG1, *Isotricha* sp YL-2021b SAG3, *Metadinium minomm* SAG1, *Ophryoscolex caudatus* SAGT3, *Ostracodinium dentatum* SAG1, *Ostracodinium gracile* SAG1 and *Polyplastron multivesiculatum* SAGT3. The genome of *Fibrobacter succinogenes* S85 (NCBI ID: 932) was also added to both rumen databases since it is well recognized as a primary cellulolytic bacterium in the rumen microbiome and has previously been observed as an active microorganism in similar studies yet was not thoroughly binned as a unique MAG in this study. Nine available fungal genomes downloaded from Joint Genome Institute (JGI) Mycocosm^55^ were added to RUDB-C (*Anaeromyces* sp. S4^25^, *Caecomyces churrovis*^*28*^, *Neocallimastix californiae*^*25*^, *Neocallimastix lanati*^*29*^, *Piromyces finnis*^*25*^, *Piromyces* sp. E2^25^, *Piromyces* UH3-1^24,25,27^, *Piromyces* eukMAG^56^, *Orpinomyces* sp.^26^). In total the complete databases consisted of 1 291 083 and 1 190 938 protein entries for RUDB-G and RUDB-C, respectively.

### Metaproteomic data generation

To 300 μL of rumen fluid sample (in total 32 rumen fluid samples originating from four cows and four goats fed four diets included in the original study^23^) 150 μL lysis buffer (30 mM DTT, 150 mM Tris-HCl (pH=8), 0.3% Triton X-100, 12% SDS) was added together with 4 mm glass beads (≤ 160 μm), followed by brief vortexing and resting on ice for 30 mins. Lysis was performed using FastPrep-24 Classic Grinder (MP Biomedical, Ohio, USA) for 3 × 60 seconds at 4.0 meter/second^57^. Samples were centrifuged at 16 000 × *g* for 15 minutes at 4°C and lysate was carefully removed. Protein concentration was measured using the Bio-Rad DC™ Protein Assay (Bio-Rad, California USA) with bovine serum albumin as standard. Absorbance of sample lysates was measured at A750 on BioTek Synergy H4 Hybrid Microplate Reader (Thermo Fisher Scientific Inc., Massaschusetts, USA). 40-50 μg of protein was prepared in SDS-buffer, heated in water bath for 5 minutes at 99°C and analysed by SDS-PAGE using Any-kD Mini-PROTEAN TGX Stain-Free gels (Bio-Rad, California, USA) in a 2-minute run for sample clean-up purposes, before staining with Coomassie Blue R-250. The visible bands were carefully excised from the gel and divided as 1×1 mm pieces before reduction, alkylation, and digestion with trypsin. Peptides were concentrated and eluted using C18 ZipTips (Merck Millipore, Darmstadt, Germany) according to manufacturer’s instructions, dried, and analysed by nano-LC-MS/MS using a Q-Exactive hybrid quadrapole Orbitrap MS (Thermo Fisher Scientific Inc., Massaschusetts, USA) as previously described^58^

### Metaproteomic data analysis

Mass spectrometry (MS) raw data were analysed with FragPipe version 19 and searched against the sample-specific protein sequence database (1 291 083 and 1 190 938 protein entries for RUDB-G and RUDB-C, respectively) with MSFragger^59^ (**Supplementary Table S4**). The database was supplemented with contaminant protein entries, such as human keratin, trypsin, and bovine serum albumin, in addition to reversed sequences of all protein entries for estimation of false discovery rates (FDR)^60^. Oxidation of methionine and protein N-terminal acetylation were used as variable modifications, while carbomidomethylation of cysteine residues were used as fixed modification. Trypsin was chosen as digestive enzyme, maximum missed cleavages allowed was one and matching tolerance levels for both MS and MS/MS were 20 ppm. The results were filtered to 1% FDR and quantification was done using IonQuant^61^ including normalization between samples and the feature ‘match between runs’ to maximize protein identifications. Perseus^62^ version 1.6.2.3 was used for further analysis. A protein group was considered valid if it was quantified in at least 50% of the replicates in at least one condition. Protein groups identified as potential contaminants were removed. Calculations of individual genome/SAG/MAG contributions were calculated using the workflow outlined by Kleiner et al.^63^, which sums protein abundances (label free quantification (LFQ) values) for all proteins assigned to each genome/SAG/MAG, and differential abundance between diets were detected by an unpaired two-sided Student’s t-test (*p-*value <0.05).

After filtration, we resolved 3657 unique protein groups across the 16 samples from cattle and 3818 unique protein groups across 15 samples from goats (**Supplementary Table S5** and **Supplementary Table S6**). Dot plots for **Figure 1** were made with ggplot2^64^ in R (v. 4.2.0)^65^. To determine which expressed metabolic pathways *En. caudatum* were significantly enriched for in each diet/animal, we used the hypeR() function from the hyperR package^66^ in R which employs the hypergeometric test for significance. The ‘geneset’ for hyperR was generated by using the keggGet() functionof KEGGREST R package to retrieve entries from the KEGG database and determine which pathways the *En. caudatum* KOs belong to. The geneset was then manually curated to only include metabolic pathways of interest (i.e., we remove pathways such as “Huntington disease”). For the ‘background’ setting in hyperR(), to be conservative, we used the total number of unique KOs (7592) in the *En. caudatum* genome that could possibly be expressed. Significantly enriched KOs with *p*-value < 0.05 were manually filtered. Both *p*-values and FDR-adjusted *p-*values are available in **Supplementary Table S7**, with FDR-adjusted *p-*values < 0.05 highlighted in bold.

**Figure 1.**
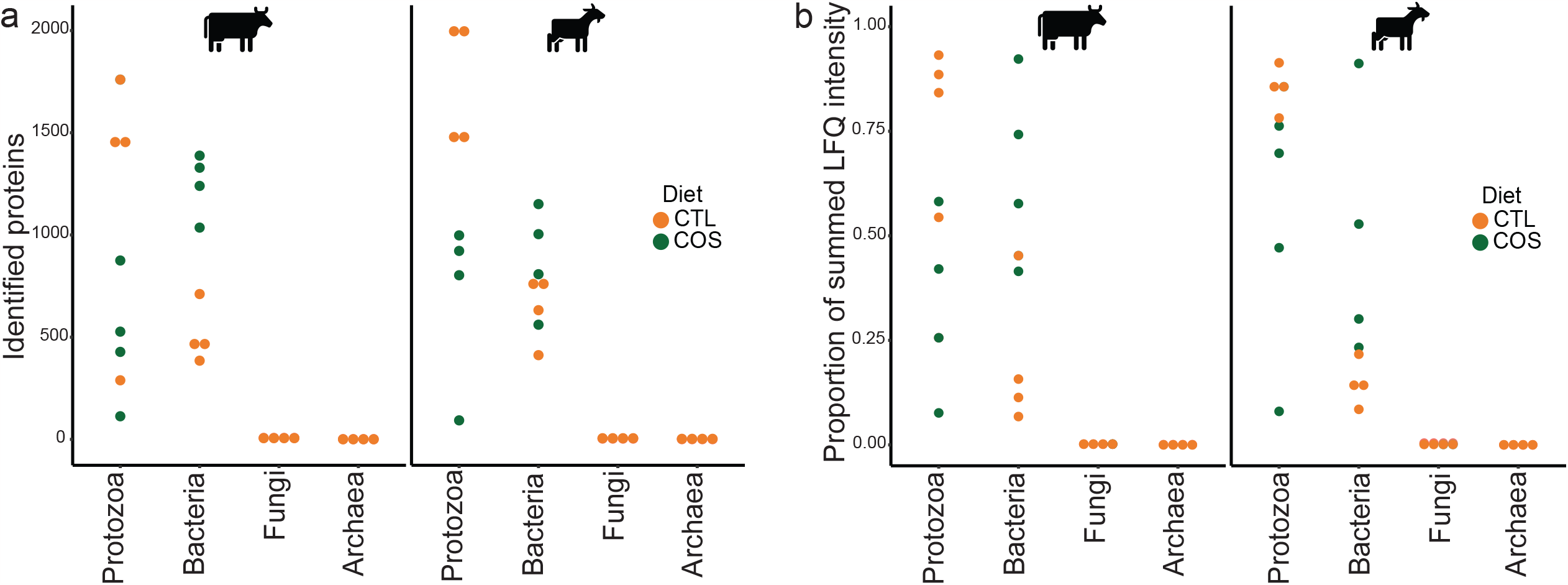
Quantities of identified protozoal proteins in the rumen microbiome vary depending on host animal and dietary conditions. Dot plots for total proteins identified (**a**), and average recovered metaproteomic expression (**b:** presented as proportion of summed LFQ intensities) belonging to protozoal, bacterial, archaeal, or fungal species across the control diet (CTL) and diets supplemented with corn oil and wheat starch (COS) for dairy cattle (n=4) and goats (n=4). Detected protein abundances for protozoal and bacterial populations can be found in **Supplementary Table S6**.

### Animal trial, sample handling and metagenomic data generation for independent validating dataset

Samples were also analysed from previously performed feeding experiments with Holstein Friesian bulls^67^. In brief, these bulls were subjected to either *ad libitum* or restricted feeding regime in a compensatory growth model detailed in Keogh *et al*., 2015^67^. Both feeding groups received the same ratio of concentrate and grass silage, respectively 70% and 30%, of which the concentrate was mainly composed of starch-rich rolled barley (72.5%) and soya (22.5%). Rumen samples were collected at slaughter and stored at -80C prior to metagenomics and metaproteomic analysis in this study.

Sample preparation, cell lysis and extraction of DNA was carried out as previously described by McCabe *et al*.^68^ Quality check of fastq files and removal of low-quality reads was performed using fastp (V.0.19.5). Sequence reads were mapped against the bovine genome (ARS-UCD1.3) using minimap2 (V.2.16), and host sequences were removed. Reads were co-assembled using Megahit (V1.2.6) with “−meta-large” pre-set option as the metagenome was complex. Metagenomic binning was applied to the co-assembly using MetaBAT2 using standard parameters (V.2.12.1). MAGs were then dereplicated using dRep (V.1.4.3), and the resulting MAGs were taxonomically assigned using Bin Annotation Tool (BAT), available on (https://github.com/dutilh/CAT). This tool internally uses prodigal (V.2.6.3) for gene prediction and DIAMOND (V.0.9.14) for the alignment against the non-redundant (nr) protein database (as of Feb 2020).

Sample preparation for metaproteomics was done by lysing cells with bead beating with two glass bead sizes (≤106 μm and 0.5 mm), in 100 mM Tris, pH8, 5% SDS and 10 mM DTT. A FastPrep 24 instrument was operated for 3 × 45 seconds at a speed of 6.5 m/s. The samples were centrifuged for 15 minutes at 20.000 × *g* and the protein extracts were cleaned by Wessel-Flügge precipitation^69^; pellets were dissolved in 5% SDS, 100 mM Tris-Cl, pH8, 10 mM DTT and kept at -20 °C until further processing. Protein digestion was performed using suspension trapping (STrap)^70^, dried in a SpeedVac (Eppendorf Concentrator Plus) and re-dissolved in 0.05 % trifluoroacetic acid, 2% acetonitrile for peptide concentration estimation using a Nanodrop One instrument, and subsequent MS/MS-analysis. The samples were analyzed using an Ultimate3000 RSLCnano UHPLC coupled to a QExactive hybrid quadrupole-orbitrap mass spectrometer (Thermo Fisher, Bremen, Germany) as described previously^58^. MS raw data were analysed with FragPipe and searched against the Holstein Friesian bulls sample-specific protein sequence database (2 533 362 protein entries) using the same workflow described above.

## RESULTS AND DISCUSSION

### Protozoal populations have large proteomes in the rumen microbiome

Because of their large size protozoal species can comprise a significant fraction of the microbial biomass in the rumen^9^. While the total count and diversity of protozoal species are lesser than their bacterial counterparts in the rumen, their genome size and total number of genes encoded within are considerably larger. In addition, due to alternative splicing and post-translational modifications, the protein representation, or proteome, of protozoal populations will be larger than the number of genes in their protozoal genome^71^. Thus, the amount of protein in a protozoa species can be expected to far exceed the amount of protein in a bacterial species that can be identified and quantified in proteomic studies. In this context, our metaproteomic data showed an extensive fraction of detectable proteins affiliated to protozoal species, and other closely related species, in both dairy cattle and goats in proportion to the combined bacterial species that were represented in our genome databases (**Figure 1a**). Protein detection intensity of proteins assigned to protozoa were also proportionally greater than that of the bacterial fraction of the rumen microbiome in animals fed the control (CTL) diet (**Figure 1b**) further supporting the significant levels of protozoal detection in our samples. Proteomes from entodiniomorph species were detected at high levels in both animal hosts, with *Ep. cattanei* and *En. caudatum* dominating the rumen metaproteome of cattle and goats respectively (**Figure 2**). In cattle we also detected entodinomorphs *D. dentatum, En. bursa* and *Ep. caudatum* (**Figure 2a, Supplementary Table S6**) at high levels whereas in goats *Entodinium* species *En. caudatum, En. longinucleatum* and *En. bursa* where among the highest detected entodinomorphs (**Figure 2c**). More than three times as many proteins from *En. caudatum* were detected in goats than cattle, however this was somewhat expected given the 7x higher counts of entodiniomorph concentration (cells/mL) previously observed in the goat samples compared to cattle (**Table 1**)^5^. In agreement with microscopic cell counts of holotrichs, we detected proteins from both *Dasytricha* spp. and *Isotricha* spp (**Table 1**) in both cattle and goats (**Figure 2a and 2c**).

**Figure 2.**
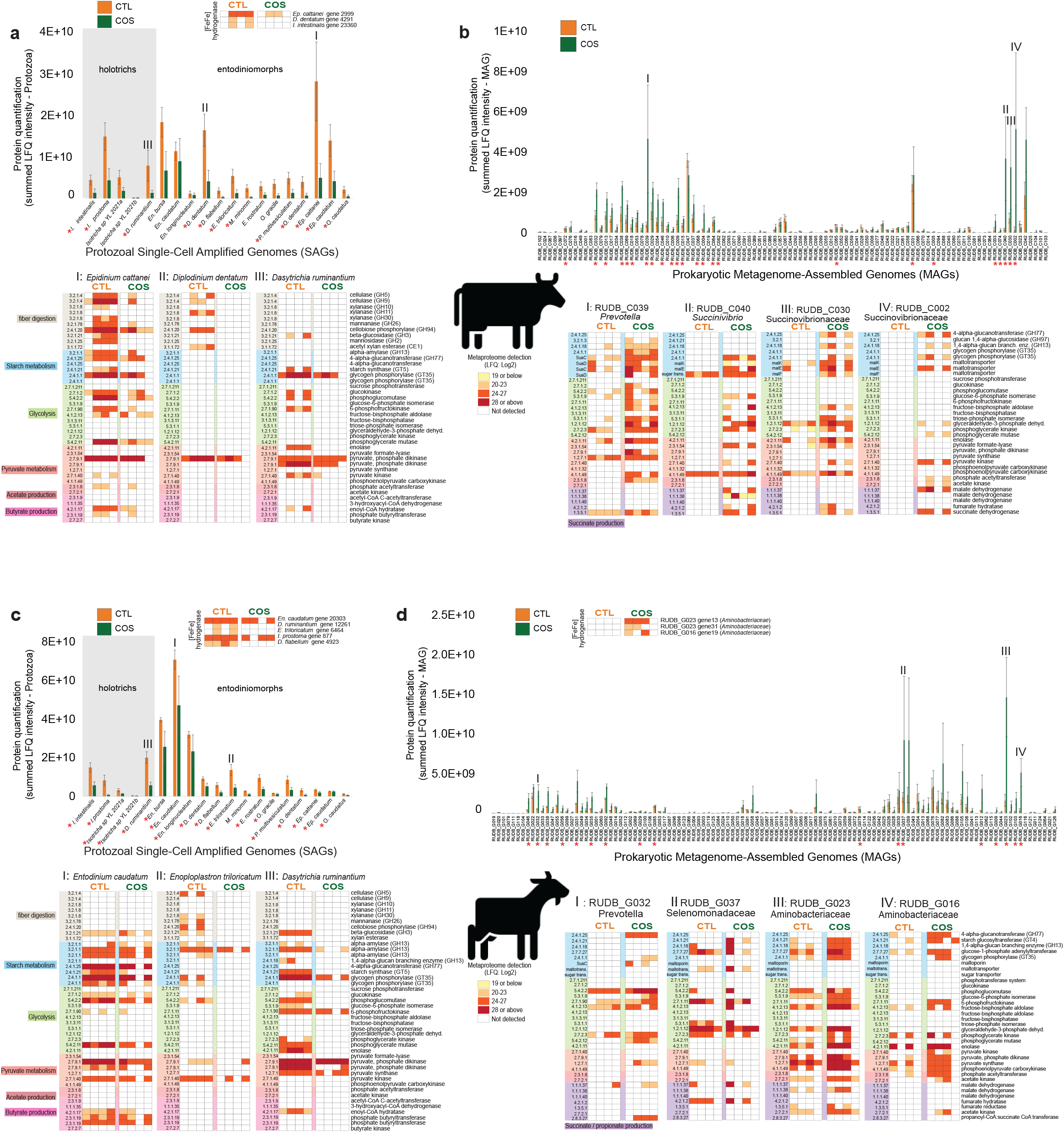
Detected proteins mapped to protozoal genomes/SAGs and bacterial metagenome-assembled genomes (MAGs) in the rumen microbiome of dairy cattle (n=4) and goats (n=4) fed either a control diet (CTL) or one supplemented with corn oil and wheat starch (COS). The figure displays metabolically active populations (as genomes, SAGs or MAGs), with selected expressed proteins (presented as Enzyme Commission (EC) number with short enzyme descriptions) active in starch degradation, glycolysis and production of pyruvate, butyrate, acetate, and succinate in cattle (**a-b**) and goats (**c-d**) fed CTL or COS diets. Panels **a** and **c** depict protozoal proteomes that were detected in cattle and goats respectively are presented separately to bacteria (panels **b** and **d**) as the scale of their protein quantification values were ∼10x larger. Protein quantification values (y-axis) were calculated by considering both the number of proteins detected per MAG/SAG/genome and their LFQ intensity: we averaged LFQ intensities for each detected protein across biological replicates for each dietary condition (CTL: green or COS: orange), which were subsequently summed for all detected proteins per MAG/SAG/genome. Heatmaps show selected MAG/SAG/genome with metabolically active proteins, presented as EC numbers, recovered from cattle (RUDB-C) and goats (RUDB-G) fed either CTL or COS diets. Where average protein abundance level for a given MAG/SAG/genome was significantly different (determined by paired Wilcoxon test, *p-*value < 0.05) between the COS and CTL diet, the given MAG/SAG/genome has been marked with a red star. MAGs are presented with their MAG ID and taxonomic annotation from GTDB-tk. Genome annotations and LFQ intensities used to create heatmaps can be found in **Supplementary Table S6**.

### Entodiniomorphs and holotrichs have influential roles in rumen carbohydrate metabolism

Rumen protozoa have long been characterized as degraders of plant fibers as well as non-structural storage polysaccharides such as starch^72^. Closer examination of the proteomes from both highly detected *Epidinium species Ep. cattanei* and *Ep. caudatum* in cattle highlighted a plethora of detected carbohydrate active enzymes (CAZymes) targeting lignocellulosic polysaccharides such as cellulose (GH5, GH9), xylan (GH10, GH11, GH30, CE1), mannan (GH26) and fiber-derived oligosaccharides (GH2, GH3, GH94) (**Figure 2a, Supplementary Table S6**). In goats similar CAZy families were detected but from different entodinomorphs species, such as *E. triloricatum* and *P. multivesiculatum* (**Figure 2c, Supplementary Table S6**). Our *in vivo* CAZyme observations were in strong agreement to long standing *in vitro* data, which have shown entodinomorphs such as *Epidinium, Polyplastron* and *Enoploplastron* have greater endoglucanase and xylanase activity, while *Entodinium* spp. have only weak activity^72^. Similarly, it has previously been shown *in vitro* that holotrichs such as *Dasytricha* spp. have glucosidase and cellobiosidase activity but negligible fibrolytic activity^73^, which our *in vivo* data also supported (**Figure 2a and 2c**). The composition of protozoal species metabolising starch (both degrading and synthesizing) also varied between the two animal hosts with goats dominated by *Entodinium* spp. And *D. ruminantium* while protozoal starch metabolism in cattle was *Epidinium* spp., *D. ruminantium and Isotricha* spp (**Figure 2, Supplementary Table S6**). While bacterial CAZymes targeting lignocellulose were also detected (e.g. GH48 cellulases from *Ruminococcaceae* populations in cattle), CAZymes targeting lignocellulose and starch were detected at higher frequency and protein quantification levels in protozoal species suggesting they have a central role in ruminal fibre digestion.

### Metabolism of En. caudatum shows predatory activity and metabolism of VFAs

*En. caudatum* is renowned for its predatory activity and is acknowledged as the most abundant protozoa in the rumen, whereby it has been estimated that 0.1% of rumen prokaryotes are digested by the rumen protozoal population every minute^74^. The dominance of *En. caudatum* in our data presented an opportunity to better understand the metabolic influence of this universal species. While previous efforts have investigated the genome and transcriptome of *En. caudatum* grown in monoculture^14,75^, our metaproteomic analysis sought to reveal *in vivo* metabolism and functions of *En. caudatum* within the rumen microbiome. In accordance with Wang *et al*.^75^, our metaproteomic analysis revealed expressed proteins attributed to metabolic pathways such as carbon metabolism, glycolysis/gluconeogenesis, starch and sucrose (and glycogen) metabolism, pyruvate metabolism, oxidative phosphorylation and production of alcohol (**Supplementary Table S6**). Wang *et al*. found that as for most rumen microbes, *En. caudatum* uses carbohydrates such as starch as its primary substrate, as well as cellulose and hemicellulose to a certain degree^75^, and their transcript analyses showed that *En. caudatum* had high levels of expression of amylases and low-level expression of hemicellulases, cellulases and pectinases. Similarly, our metaproteomic analysis reveals expression of amylases by *En. caudatum* that are predicted to enable *En. caudatum* to engulf and degrade starch granules to simpler sugars and to produce glycogen, its most important storage carbohydrate^76^. However, no detection of *En. caudatum* CAZymes related to hemicellulose or pectin were observed in any of our metaproteomes, suggesting that it is not engaging in the deconstruction of these carbohydrates at the time our samples were collected for analysis (before feeding). It should be noted that ruminal fermentation activity as well as production of VFAs and methane will be at its highest after feeding, as a result of an increased availability of fermentable substrate^77^. While sampling time can influence the recovered microbial composition and hence function, any differences in metabolic parameters or species abundance in this study is comparable across both diets given the consistent sampling times.

While monoculture cultures of *En. caudatum* have not been established to verify the VFAs it can produce, Wang *et al*. found transcripts of enzymes involved in fermentative formation of acetate and butyrate^75^. Similarly, we detected proteins inferred in metabolism of acetate, butyrate, and alcohol in *En. caudatum*. Goats fed the CTL diet had a significantly higher proportion of proteins from *En. caudatum* and concurrently had increased relative levels of acetate and butyrate compared to animals fed the corn oil and wheat starch diet (COS), which had fewer *En. caudatum* proteins and lower acetate/butyrate levels (**Figure 2** and **Table 1**). As *En. caudatum* populations were seemingly most abundant in goats fed the CTL diet, we used these metaproteomes to reconstruct metabolic features (**Figure 3**). Of the 538 *En. caudatum* proteins identified in CTL-fed goats, 244 had unique KO numbers assigned, from which KEGG Mapper reconstructions^54^ enabled functional assignment of 217 proteins to metabolic pathways. Our metabolic reconstructions showed expressed proteins involved in endocytosis, phagosome and lysosome processes for predatory activity, engulfment, and digestion of bacteria (**Figure 3, Supplementary Table S6**). For the rumen samples used in this study Martin *et al*. previously observed higher NH_3_ concentrations in goats compared to cattle and hypothesised that it might have resulted from increased bacterial protein breakdown and feed protein degradability due to higher density of entodiniomorphs known for their predatory activity^5^. In support of these observations, we performed metaproteomic pathway enrichment analysis of *En. caudatum* (**Figure 3, Supplementary Table S7**) proteins detected in goats, which revealed significantly enriched endocytosis metabolism. Other biological processes such as glycolysis and starch and sucrose metabolism were significantly enriched pathways in *En. caudatum* (**Supplementary Table S7**). Although suspected of having metabolic interactions with methanogenic archaea, several protozoal populations such as *Entodinium* are hypothesized as having associations with certain members of the Gram-negative *Gammaproteobacteria*, which multiple studies have speculated are resistant to protozoal engulfment^78-80^. In contrast, Gutierrez and Davis previously demonstrated that *Entodinium*-species engulf Gram positive starch-degraders^80^. In the context of our data, we speculate that CTL fed animals provided *En. caudatum*-like populations optimal conditions for predation, whereas increased starch levels in the COS diets facilitated increases in Gram-negative species that are possibly resistant to protozoal engulfment and/or reduced pH levels, leading to sub-optimal conditions for *Entodinium* and a decrease in its levels (**Figure 2**).

**Figure 3.**
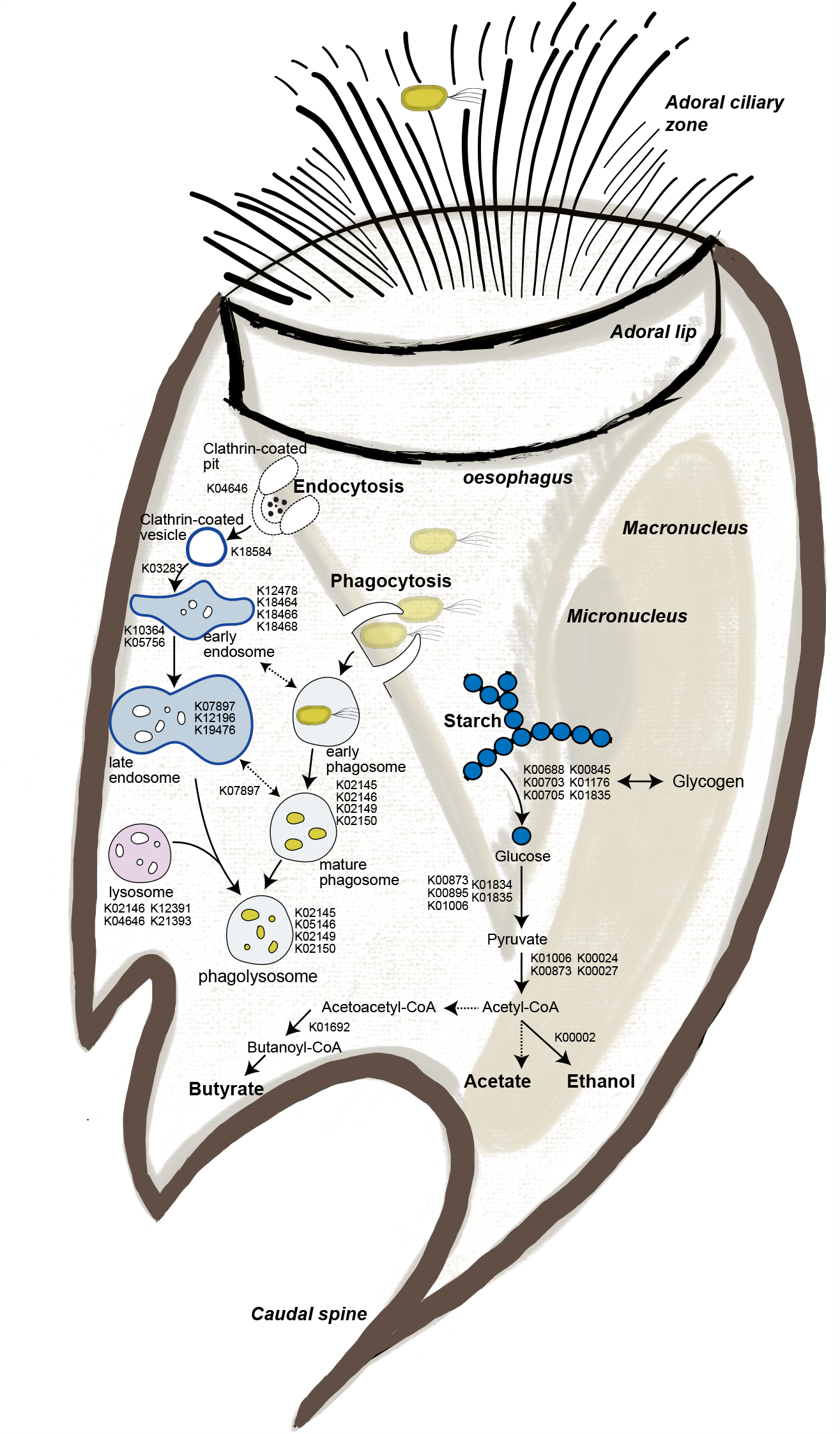
Reconstructed phagolysosome formation and starch metabolism of *Entodinium caudatum* within the rumen microbiome of goats fed the control diet (CTL) based on metaproteomic analysis. KO identifiers for identified proteins were analysed via KEGG mapper to reconstruct expressed key features in the metabolism of *En. caudatum*. Dashed arrows represent proteins or pathways that were not detected in our metaproteomes but are key steps in their respective pathways. Detailed information connecting KO identifiers to their respective gene ID, LFQ and animal/diet can be found in **Supplementary Table S6**.

### Protozoa were less active in diets supplemented with starch regardless of their starch metabolising reputations

The changes in VFA and methane levels in animals fed the high starch COS diet, previously measured by Martin *et al*.^5^, suggested significant alterations in composition and thus functions of the rumen microbiome irrespective of host species. In particular, a decrease in proportions of acetate and butyrate, decrease in the acetate:propionate ratio and an increase in relative propionate levels were observed in animals fed the COS diet, compared to the CTL diet (**Table 1**). Diets that are high in starch content or with low forage:concentrate ratios have previously been shown to result in higher production of propionate and succinate, as they are easily fermented in the rumen and accordingly have high passage rates^81,82^. We therefore leveraged our genome-centric metaproteomic data from both cattle (**Figure 2a-b)** and goats (**Figure 2c-d**) fed either the COS or CTL diet to gain an overview of protein expression from individual populations. We specifically focused on pathways involved in the degradation of starch (CTL: corn starch, COS: corn + wheat starch) to pyruvate through glycolysis and finally formation of acetate, butyrate, and propionate (via succinate). Irrespective of host, and despite their starch-metabolising reputation^75,76^, *Entodinium, Epidinium, Isotricha* and *Dasytricha* spp had lower abundance and less proteins involved in starch degradation in animals fed the COS diet compared to those fed the CTL diet (**Figure 2a** and **2c**). Further, we observed opposing patterns for proteins identified as *Entodinium* and *Epidinium*-spp. involved in glycolysis, and production of pyruvate, acetate, and butyrate, which were detected in higher levels in both cattle and goats fed the CTL diet compared to the COS diet.

While several putative protozoal amylases were detected across all animals and diets, their quantification levels (i.e., protein abundances) did not increase as expected when higher levels of starch were available (**Figure 2a** and **2c**). We therefore hypothesized that the observed shift in VFA profiles in response to increased starch was additionally influenced by the bacterial fraction of the rumen microbiome. In contrast to lower levels of protozoa in the animals fed the COS diet, we observed an increase in bacterial proteome detection irrespective of host (**Figure 4**), including suspected starch-degrading and succinate- and propionate-producing bacterial species (**Figure 2b** and **2d**). For example, starch fermentation pathways from population genomes affiliated with the *Succinivibrionaceae* family, *Prevotella* species and, additionally for goats, members of the *Aminobacteriaceae* families, were detected at higher proteomic levels in the animals fed the COS diet compared to those fed the CTL diet (**Figure 2b** and **2d**).

**Figure 4.**
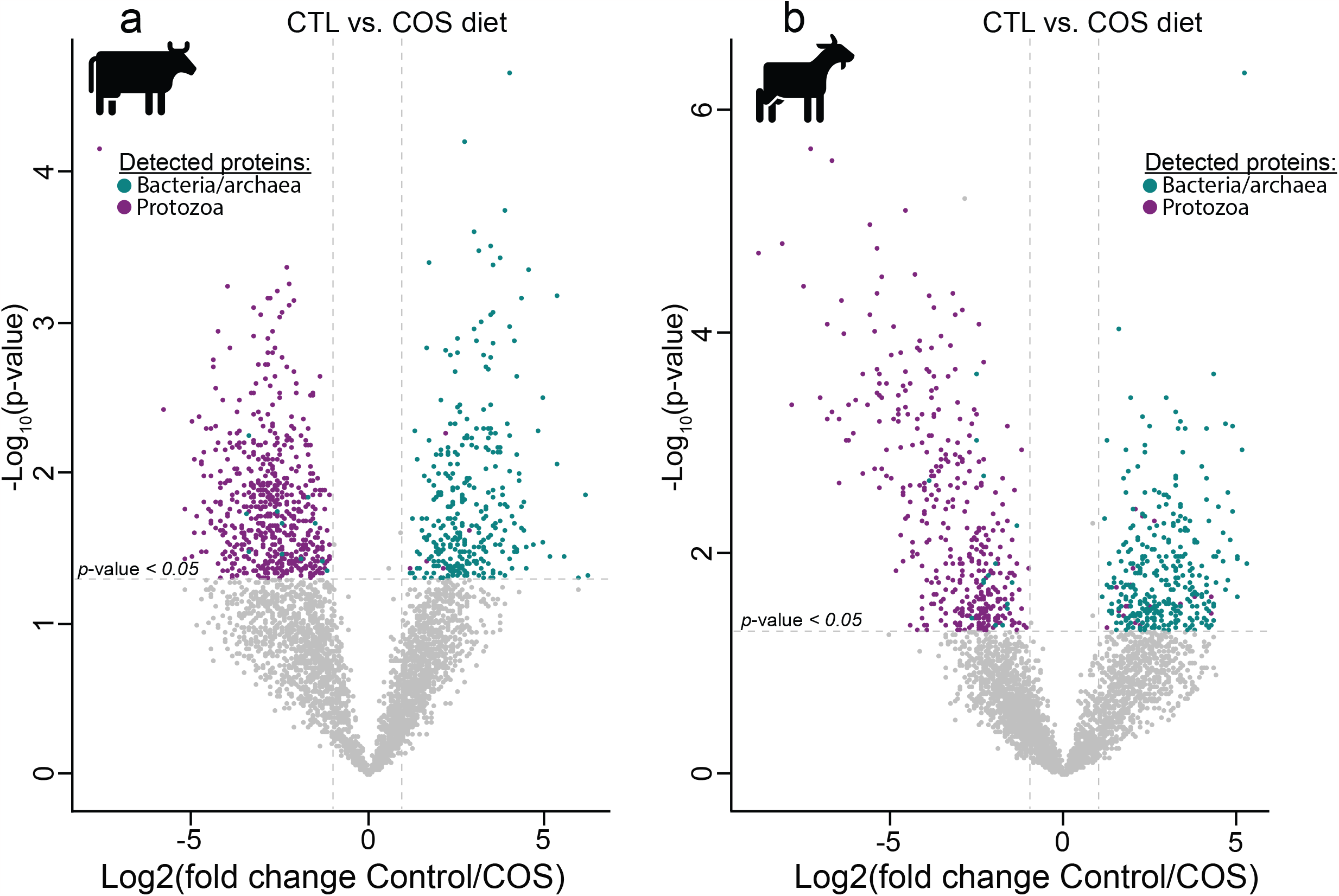
Volcano plots indicating different rumen microbiome proteins from dairy cattle (a) and goats (b) fed either the control or COS diets and which displayed both large magnitude of fold-changes in LFQ intensities (x axis) and high statistical significance (-log10 of nominal *p-*values using an unpaired t-test, y axis). Dashed horizontal line denotes a *p-*value <0.05 cut-off. For both animal hosts we observed an increase in protein detection for bacterial populations (greenblue) in animals fed high-starch diets compared to protozoal populations (purple), which were detected at higher LFQ intensities in animals fed the control diet.

### Protozoa are less active in animals that produce lower methane yield

For the animals sampled in this study, Martin *et al*. demonstrated a ∼25-30% reduction in methane emissions in both cattle and goats fed the COS diet compared to the control (**Table 1**)^5^. Previous comparisons between defaunated and faunated animals have shown decrease in methane production in protozoa-free ruminants, suggesting symbiotic interactions between methanogenic archaea and protozoal species^78^. Methanogen’s epi- and endo-symbiotic relationships with protozoa have also been suggested to contribute to 9-37% of rumen methanogenesis^78,83-85^. Moreover, studying microcosms with the presence and absence of protozoal species Solomon *et al*. reported higher levels of acetate and butyrate in microcosms with protozoa present in addition to increased methane emissions^78^, which supports the main findings of animals fed the CTL diet in this study (**Table 1**). Several of the protozoal species detected in our metaproteomic data are known to have hydrogenosomes and closer examination showed that [FeFe]-hydrogenases from both holotrich and entodinomorph genera, such as *Dasytricha, Isotricha, Diplodinium, Epidinium, Entodinium* and *Enoploplastron*, were detected at higher quantification levels in animals fed the CTL compared to those fed COS (**Figure 2a and 2c, Supplementary Table S6**). We also observed partial evidence in our proteomic data that the hydrogenosome-lacking *En. caudatum* makes contributions to ruminal hydrogen production, supporting previous reports associating its abundance with higher methane levels^78,85,86^. Only one of its eight [FeFe]-hydrogenases were detected in goats (absent in cattle), though it showed no changes in protein abundance in either the high (CTL) or low (COS) methane yielding animals. Given our observations, we speculate that predicted protozoal-methanogen relationships that lead to increased methane levels in this study are centred on hydrogen transfer.

The addition of lipids in diets for ruminants have been largely studied as a methane mitigation strategy^87,88^. However, it is widely believed that lipids are not fermented in the rumen and rather modified/hydrogenated and hence do not contribute to hydrogen production or methane production in the rumen. While biohydrogenation of unsaturated fatty acids such as corn oil can serve as a hydrogen sink, it has been observed that very small amounts of metabolic hydrogen (1-2%) are used for biohydrogenation^87,88^. Several studies have also shown that the CH_4_ mitigation effects of lipids are dependent on both dose and fatty acid composition^89,90^. For example, medium chain fatty acids (MCFAs), such as lauric acid, have been proven to be a more effective inhibitor of protozoa than long chain polyunsaturated fatty acids (PUFAs), such as corn oil, which mainly consists of linoleic acid^88-91^. In a study conducted by Zhang *et al*.^90^, goats fed corn oil as a supplement decreased ruminal hydrogen concentrations and total methane emissions. Nevertheless, there was seemingly no effect on rumen protozoal populations, which suggests that the corn oil dosage used in the Zhang study did not act as an anti-protozoal agent^90^. In addition, high levels of PUFAs in the diet will also likely impact the metabolism of various bacterial and archaeal species^88-90^, particularly keystone fibre degrading Gram-positive bacterial species that produce various levels of VFAs and hydrogen^92,93^. Collectively, these previous findings were in agreement with the decreased CH_4_ production in cattle and goats in this study fed the COS diet, which was observed to additionally impact other bacterial metabolism and ruminal fermentation parameters, such as increased propionate and decreased butyrate and acetate levels (**Table 1**)^5^. Thus, while we cannot unequivocally rule out that diets supplemented with lipids at higher levels could be directly affecting the abundance of protozoa, we believe it is also possible that corn oil is influencing other fibre-degrading and hydrogen-producing species in the rumen, hence effecting ruminal fermentation and methane emissions^88,94^. Nonetheless, validation of these results under different conditions and extended sampling is necessary to verify our findings.

Increases in dietary starch for ruminants is known to stimulate the propionate and succinate pathways of starch-degrading bacteria, which due to their net incorporation of metabolic hydrogen [H] represent a [H] sink in rumen fermentation besides hydrogenotrophic methanogenesis^83,95^. Diets rich in starch are more fermentable in the rumen, which can decrease the ruminal pH to levels that can inhibit methanogenic archaea and fibre-degrading bacterial species^96,97^. Yet, lowered pH levels in the rumen can also lead to clinical (or sub-clinical in most production scenarios) ruminal acidosis^98,99^. Hence, high concentrate diets, which increase production of propionate at the expense of methane, does not necessarily opt for a viable methane mitigation strategy in the long term. Our results suggests that decreased methanogenesis in COS-fed animals is likely due to a decrease in available hydrogen and/or decrease in pH levels, which we predict is caused by the lower levels of hydrogen-producing protozoa and bacteria as well as the metabolism of dominant bacterial populations that likely do not produce exogenous hydrogen due to their own [H]-utilizing succinate and propionate metabolism. This prediction was partially supported by the detection of several putative hydrogenases believed to use molecular hydrogen in putative fumarate-reducing *Aminobacteriaceae* populations that were dominant in the COS-fed goat rumen. In particular, two [FeFe]-hydrogenases were detected in COS-fed animals from RUDB_G023 (ORF_4456741_13) and RUDBG016 (ORF_ 1143158_19) that were annotated as NADP-reducing hydrogenase subunits (EC:1.12.1.3) (**Figure 2d, Supplementary Table S6**).

### Protozoal dominance is validated in geographically independent datasets

To further test our hypothesis that protozoal species play a central role in the rumen ecosystem, we explored additional metagenome-centric metaproteomic datasets originating from an independent feeding experiment performed in Ireland on 60 Holstein Friesian bulls^67^. In brief, these bulls were subjected to the same ratio of concentrate and grass silage at either an *ad libitum* or restricted feeding regime in a compensatory growth model detailed in Keogh et al, 2015^67^. We applied the same strategy as for the described Holstein dairy cattle and alpine goats to resolve the metaproteomic dataset for a subset of 15 animals (7 restricted and 8 *ad libitum*) against 781 reconstructed sample-specific MAGs (RUDB-HF), which were supplemented with the genome of *En. caudatum*, 18 protozoal SAGs^15^ as well as genomes of available anaerobic fungi. This collection of microbial prokaryote and eukaryote genomes was then used as a sequence database for the generated protein spectra. Consistent with our previous observation in dairy cattle, a substantial proportion of the detected proteins were affiliated to *Epidinium* (*Ep. cattanei* and *Ep. caudatum*), *Entodinium* (*En. caudatum* and *En. bursa*) and *Isotricha* spp. providing further support that protozoa are an important and metabolically active contributor to the rumen microbiome (**Figure 5, Supplementary Table S8**). The protein quantification (measured as sum of LFQ intensities affiliated to each MAG/genome, averaged for each diet) was also higher in the rumen sample from bulls on the restricted diet, which likely had less starch available compared to the *ad libitum* group and a higher retention time (**Figure 5a, Supplementary Table S8**). A previously published 16S rRNA amplicon investigation of the phylogenetic differences between the rumen microbiomes of these two diet groups highlighted an increase in *Succinivibrionaceae* in the starch-rich *ad libitum* diet^68^. Our metaproteomic analysis confirmed a significantly higher (unpaired t-test, *p-*value < 0.05) proteomic detection of several *Succinivibrionaceae*-MAGs under the *ad libitum* group (**Supplementary Table S8**), accompanied with a reduced acetate:propionate ratio in the rumen, which is often associated with increased feed efficiency and reduced production of methane^68^. These observations largely mirror the dominance of *Succinivibrionaceae*-MAGs in the dairy cattle and goats fed the COS diet, further strengthening our hypothesis that *Entodinium* and *Epidinium* spp do not metabolically respond to increases in available starch in the host animals’ diets and have other roles than being a primary starch degrader.

**Figure 5.**
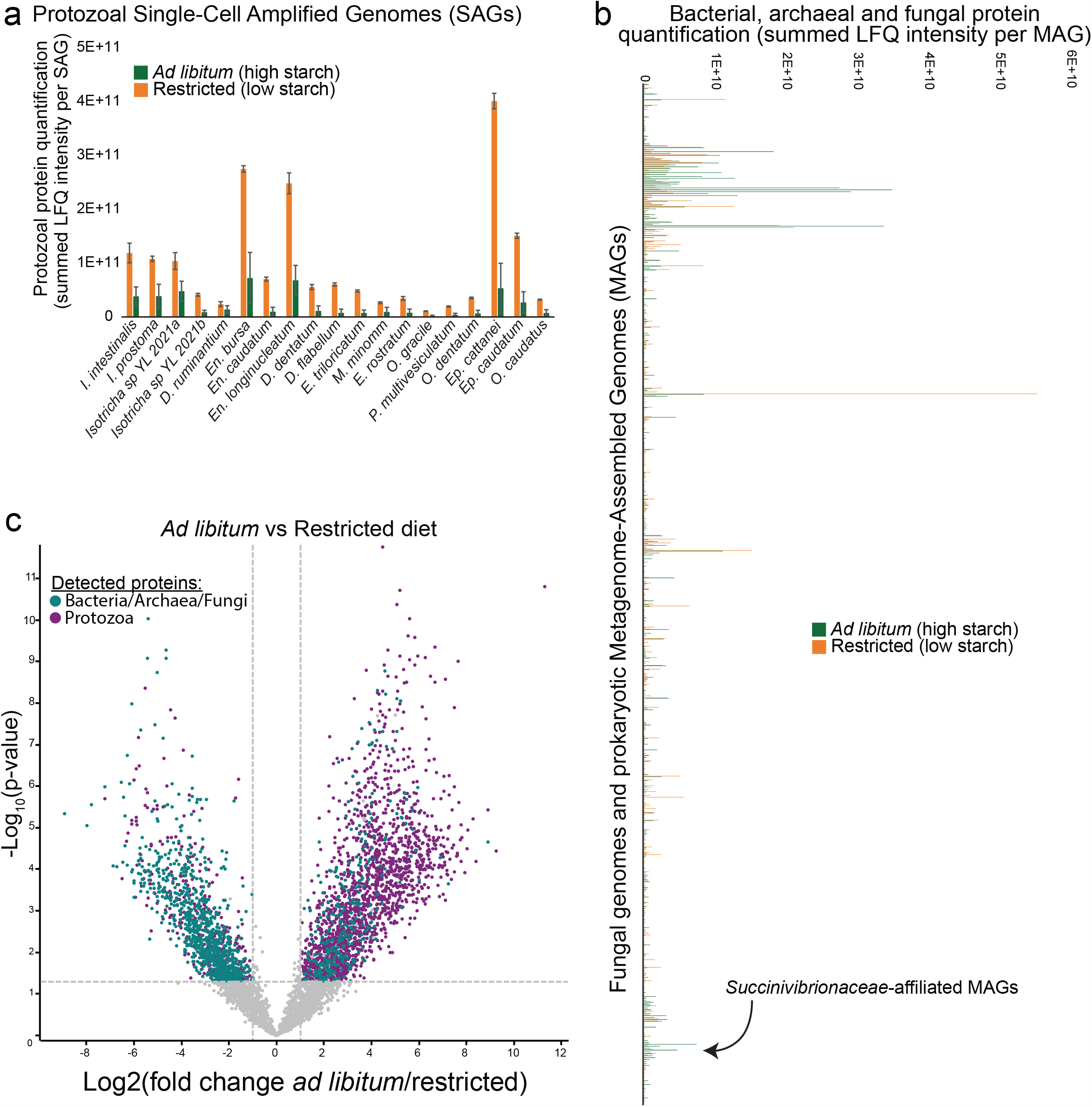
The proteomes of rumen microbiome populations from Holstein-Friesian beef cattle are affected by high starch diets. A total of 60 beef cattle were subjected to two dietary contrasting condition: 30 animals with *ad libitum* feeding and 30 subjected to 125 days of feed restriction. Dietary components in both treatments consisted of 70% concentrate, and 30% grass silage, with the concentrate containing rolled barley 72.5%, soya 22.5%, molasses 3% and calf mineral 2%. Rolled barley is high in energy and starch content (∼50%). Metaproteomes for a subset of 15 animals (7 restricted and 8 *ad libitum*) were analysed against a database that combined 19 protozoal SAGs/genome (**a**) and 781 MAGs and isolate genomes (**b**), including fungal representatives. Protozoal proteomes are presented separately as the scale of their protein quantification values were ∼10x larger. Protein quantification values were calculated by considering both the number of proteins detected per MAG/SAG/genome and their abundance; we averaged LFQ intensities for each detected protein across biological replicates for each dietary condition (*ad libitum*: green or restricted: orange), which were subsequently summed for all detected proteins per MAG/SAG/genome. Similar to our observations in Holstein dairy heifers and Alpine goats (**Figure 2)**, the proteomes of protozoal species were a major fraction of the total rumen metaproteome, however they were substantially reduced in dietary conditions where starch content was higher. Further supporting our results in dairy cattle, we observed in high-starch conditions an increase in protein detection for bacterial populations affiliated to the *Succinivibrionaceae*. **c**. Volcano plot indicating different rumen microbiome proteins that displayed both large magnitude of fold-changes in LFQ intensities (x axis) and high statistical significance (-log10 of nominal *p-*values using an unpaired t-test, y axis). Dashed horizontal line denotes a *p*-value<0.05 cut-off. Volcano plots showed an increase in protein detection for bacterial populations (greenblue) in animals fed *ad libitum* compared to protozoal populations (purple), which were detected at higher LFQ intensities in animals under restricted conditions. LFQ intensities used to create panel **a** and **b** can be found in **Supplementary Table S8**.

To assess whether similar patterns of diet-induced microbiome shifts were consistent across the two geographically distinct datasets used in this study, we compared ratios of protein abundance level for commonly observed taxonomic families. Spearman’s rank correlation coefficients were calculated for the comparisons, where two tests were performed: correlation between common taxonomic families (rarely observed families with < 2 MAGs with detected proteins across both conditions were excluded) in the rumen of beef (RUDB_HF) and dairy (RUDB_C) cattle as well as beef cattle (RUDB_HF) and alpine goats (RUDB_G), respectively. Changes in protein abundance levels from a total of 16 taxonomic families were found in common between the rumen microbiomes of beef and dairy cattle (significant correlation: *p-*value < 0.05 and correlation coefficient rho of 0.68), further strengthening our hypothesis that similar structural rumen microbiome shifts occur across independent datasets of cattle subjected to diets of high and low starch (**Figure S1, Supplementary Table S9**). However, unsurprisingly no significant correlations were observed between the rumen microbiomes of goats (RUDB_G) and beef cattle (RUDB_HF) (*p-*value > 0.05 and correlation coefficient rho of 0.19, **Supplementary Table S9, Supplementary Figure S1**), despite similar trends in protein abundance levels across taxa being observed (**Figure 2, Figure 5**, and **Supplementary Figure S1)**.

In conclusion, by using a (meta)genome-centric metaproteomics approach we primarily investigated the role of the rumen protozoa in the rumen microbiome of beef and dairy cattle as well as dairy goats that were subjected to varying dietary conditions. We showed that the proteomes of core entodinomorph and holotrich genera such as *Entodinium, Epidinium, Dasytricha* and *Isotricha* constitute a substantial fraction of the recovered rumen microbial proteome, which supports previous 16S/18S rRNA gene-based rumen census data that have highlighted their global dominance across a plethora of ruminant species. In both animal hosts, protozoal CAZymes targeting lignocellulose were detected though more frequently and at higher quantification levels in animals fed the CTL diet compared to the COS supplemented diet, presumably due to lower cell density in the later (**Table 1**). Proteins identified as *En. caudatum, Epidinium, Isotricha* and *Dasytricha* spp. were surprisingly detected at lower levels in animals that were fed increased levels of wheat starch, despite their reputable starch-metabolising capabilities (**Figures 2-5**). We hypothesize that protozoa were being out competed by Gram-negative bacterial species (e.g. *Succinivibrionaceae* in cattle and/or *Aminobacteriaceae* in goats) in the COS-fed animals, which were possibly resistant to protozoal engulfment^78-80^ and/or lower pH levels, creating sub-optimal conditions for starch-degrading protozoal populations. We also observed lower detection of protozoal [FeFe]-hydrogenases in COS-fed animals that measured a ∼25-30% lower CH_4_ yield at the time of sampling in this study (prior to feeding), further suggesting the specific conditions that enable succinate-and propionate-producing populations to flourish subsequently exert an impact on hydrogen and methane metabolisms in the rumen microbiome. For animals that were additionally fed corn oil, it is acknowledged that diets supplemented with lipids can have effects on the decreased abundance of protozoa, however our study did not provide evidence of a direct effect. While much work is still needed to confirm our abovementioned hypotheses, our integrated metaproteomics approaches have demonstrated the future importance of including eukaryote populations for accurate and meaningful analyses of the rumen microbiome and its impact on GHG mitigation strategies and host productivity traits.

## Acknowledgements

PBP and TOA are grateful for support from The Research Council of Norway (FRIPRO program, PBP: 250479), as well as the European Research Commission Starting Grant Fellowship (awarded to PBP: 336355 - MicroDE), and the Novo Nordisk Foundation (awarded to PBP: 0054575 - SuPAcow). LHH was supported by The Research Council of Norway (FRIPRO program, LHH: 302639 – SeaCow), while MØA was supported by the Novo Nordisk Foundation, project No. NNF20OC006131. The experimental trial was financed by APIS-GENE (Paris, France) as part of the NutriLip project. Gas emission measurements were funded by UMR 1213 Herbivores (INRAE, Saint-Genès-Champanelle, France). The sequencing service was provided by the Norwegian Sequencing Centre (www.sequencing.uio.no), a national technology platform hosted by the University of Oslo and supported by the “Functional Genomics” and “Infrastructure” programs of the Research Council of Norway and the Southeastern Regional Health Authorities. The authors acknowledge the Orion High Performance Computing Center at the Norwegian University of Life Sciences and Sigma2 - the National Infrastructure for High Performance Computing and Data Storage in Norway for providing computational resources that have contributed to meta-omics computations reported in this paper. Mass spectrometry-based proteomic analyses were performed by The MS and Proteomics Core Facility, Norwegian University of Life Sciences (NMBU). This facility is a member of the National Network of Advanced Proteomics Infrastructure (NAPI), which is funded by the Research Council of Norway INFRASTRUKTUR-program (project number: 295910).

The authors declare no conflicts of interest.

## Data Availability

Raw shotgun metagenomic data has been deposited in the National Center for Biotechnology Sequence Read Archive (NCBI-SRA) under accessions numbers SRR19524239 to SRR19524270 with links to BioProject accession number PRJNA844951. All annotated prokaryote MAGs are available publicly at DOI: 10.6084/m9.figshare.20066972.v1. The mass spectrometry proteomics data have been deposited to the ProteomeXchange Consortium via the PRIDE^100^ partner repository with the dataset identifiers PXD040467, PXD040454 and PXD040349. NCBI accession numbers for the protozal SAGs used in this study are as follows: *Dasytricha ruminantium* SAG3 (JAJJKK000000000), *Diplodinium dentatum* SAGT1 (JAJJLX000000000), *Diplodinium flabellum* SAG1 (JAJJLB000000000), *Enoploplastron triloricatum* SAGT1 (JAJNAB000000000), *Entodinium bursa* SAG3 (JAJJKR000000000), *Epidinium cattanei* SAG3 (JAJJKG000000000), *Epidinium caudatum* SAG1 (JAJMZR000000000), *Eremoplastron rostratum* SAG2 (JAJMZO000000000), *Isotricha intestinalis* SAGT2 (JAJJLT000000000), *Isotricha prostoma* SAG3 (JAJMZS000000000), *Isotricha* sp. YL-2021a SAG1 (JAJJLH000000000), *Isotricha* sp. YL-2021b SAG3 (JAJJKW000000000), *Metadinium minomm* SAG1 (JAJMZT000000000), *Ophryoscolex caudatus* SAGT3 (JAJJLZ000000000), *Ostracodinium dentatum* SAG1 (JAJJKF000000000), *Ostracodinium gracile* SAG1 (JAJJKZ000000000), *Polyplastron multivesiculatum* SAGT3 (JAJJLS000000000).

## Figure legends

**Supplementary Figure S1. Protein abundance level ratios calculated for commonly observed families between animals fed high/low starch diets**. Log 2-Fold change ratios between average LFQ intensities for dairy cattle (**a**), goats (**b**) or beef cattle from an independent dataset (**c**) fed high or low starch diets. Positive ratios show higher average protein abundance level for a given taxonomic family in animals that were fed high starch diets (COS or *ad libitum*). Negative ratios show higher average protein abundance levels for a given taxonomic family in animals that were fed the low starch diets (CTL or Restricted). Rarely observed families with < 2 MAGs with detected proteins across both conditions were excluded from the analysis. All ratios, and subsequent correlation tests are available in **Supplementary Table S9**.

