## Supplementary figures and images for "Metabolic influence of core ciliates within the rumen microbiome"

### Supplementary Figure S1

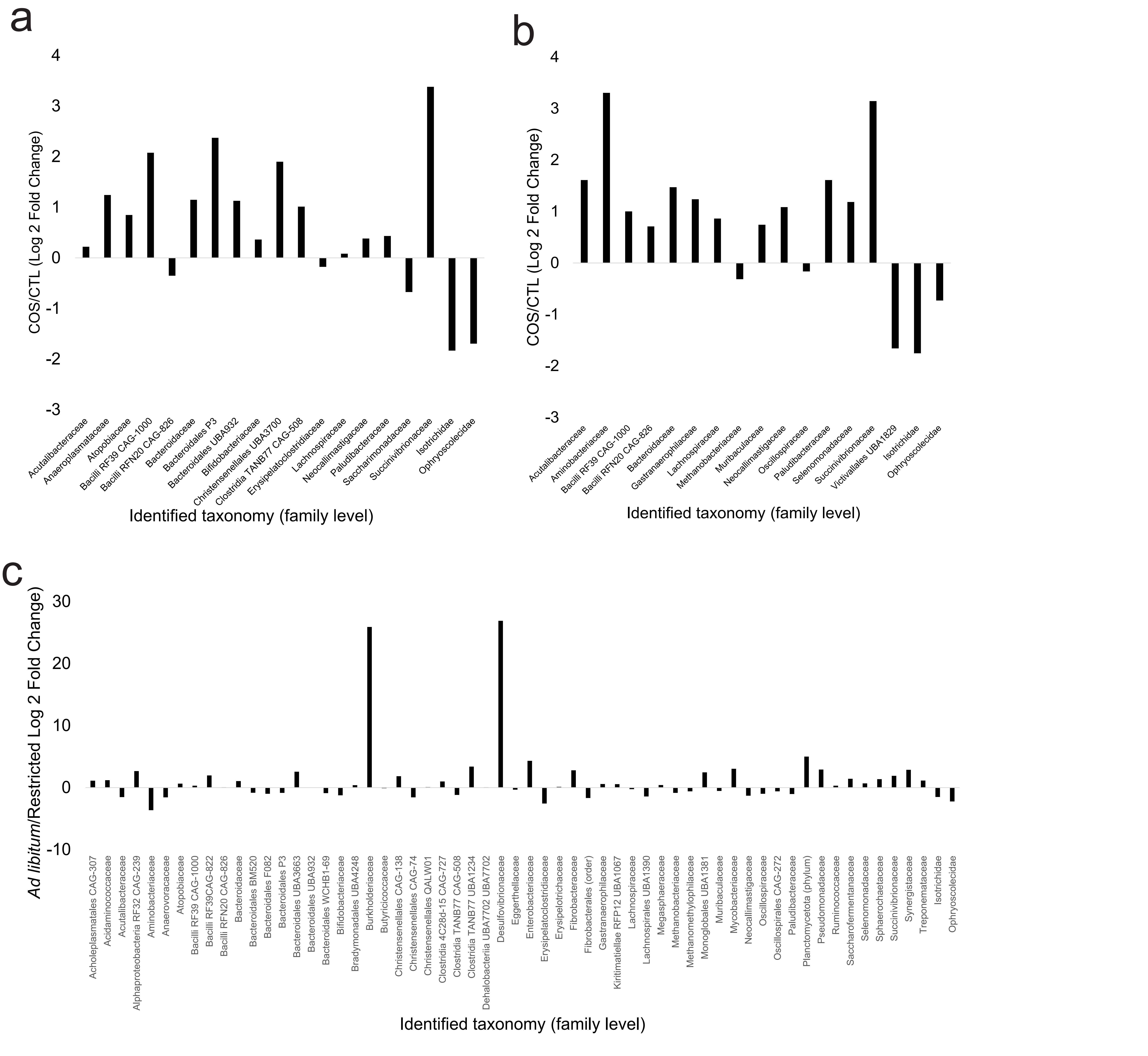
